# Comprehensive Assessment of Force-Field Performance in Molecular Dynamics Simulations of DNA/RNA Hybrid Duplexes

**DOI:** 10.1101/2024.05.06.592691

**Authors:** Barbora Knappeová, Vojtěch Mlýnský, Martin Pykal, Jiří Šponer, Pavel Banáš, Michal Otyepka, Miroslav Krepl

**Author notes:** Joint-first authors (Barbora Knappeová, Miroslav Krepl). Corresponding author: Miroslav Krepl.

## Abstract

Mixed double helices formed by RNA and DNA strands, commonly referred to as hybrid duplexes or hybrids, are essential in biological processes like transcription and reverse transcription. They are also important for their applications in CRISPR gene editing and nanotechnology. Yet, despite their significance, the hybrids have been seldom modeled by atomistic molecular dynamics methodology, and there is no benchmark study systematically assessing the force-field performance. Here, we present an extensive benchmark study of the hybrids using contemporary and commonly utilized pairwise additive and polarizable nucleic acid force fields. Our findings indicate that none of the available force-field choices accurately reproduces all the characteristic structural details of the hybrids. The AMBER force fields are unable to populate the C3′-endo (north) pucker of the DNA strand and underestimate inclination. CHARMM force field accurately describes the C3′-endo pucker and inclination but shows base pair instability. The polarizable force fields struggle with accurately reproducing the helical parameters. Some force-field combinations even demonstrate a discernible conflict between the RNA and DNA parameters. In this work, we offer a candid assessment of the force-field performance for mixed DNA/RNA duplexes. We provide guidance on selecting utilizable force-field combinations, as well as highlight potential pitfalls and best practices for obtaining optimal performance.

## Introduction

The gene expression process inevitably involves formation of mixed RNA and DNA duplexes (hybrids) during the transcription, with the newly-synthetized RNA strand temporarily base paired to its DNA template.^1^ The opposite process occurs during the reverse transcription, when the DNA is synthesized based on the RNA template.^2^ Combination of ubiquity and relatively short lifetimes makes hybrids an important but less studied component of cellular biology. The hybrids are also formed during CRISPR gene editing^3–4^ and they have a promising application in nanotechnology.^5–7^ Structurally, the hybrids form classical right-handed double helices, but the major structural biology methods somewhat disagree on the details of their geometries. The X-ray crystallography shows hybrids closely matching the A-form duplexes, including the C3′-endo pucker and χ *anti* base conformation for both strands.^8–13^ In contrast, the NMR solution experiments suggest a population split between A-and B-forms,^14–15^ pointing to a mixed population of C2′-endo and C3′-endo sugar puckers in the DNA strand of the hybrids, with concomitant χ *anti/high-anti* transitions.^16–20^ Overall, there is a notable degree of ambiguity in the experimental structural characterization of DNA/RNA hybrids, a situation which could be greatly improved by well-executed computational studies.

When performing molecular dynamics (MD) simulations of the hybrids, the choice of the nucleic acid force fields (*ff*s) is not trivial. In case of the AMBER family of *ff*s (a popular choice for simulations of nucleic acids), the *ff* parameters for DNA and RNA have been developed separately for over a decade.^21–22^ This leads to a question which combination of nucleic acid *ff*s should be utilized for hybrids to obtain optimal performance consistent with the experiments. In some cases, both RNA and DNA *ff* parameters are developed by a single research group,^22–25^ making their combined use logical. However, many more groups are separately working on deriving and optimizing parameters for either the RNA or DNA only.^26–33^ In those cases, the choice of the second nucleic acids *ff* is arbitrary and difficult to justify, especially when no specific recommendation is given by the *ff* developers and the hybrids are seldom among the systems used for benchmarking. In fact, there is currently no study systematically assessing the *ff* performance in MD simulations of hybrid duplexes available in the literature, although computational studies of specific hybrids have been successfully carried out.^34–46^

Here, we present a benchmark study that evaluates the performance of several modern AMBER nucleic acid *ff*s for MD simulations of DNA/RNA hybrids. We also test the CHARMM36^47–48^ and the latest Drude^49–51^ and Amoeba^52–53^ polarizable *ff*s. To observe the helical form changes within the same sequence context, we utilize the structure of the Dickerson-Drew B-DNA dodecamer^54^ and its modeled A-RNA and hybrid sequence equivalents. As an experimental reference for the hybrid structure, we use the polypurine tract (PPT) hybrid duplex,^13^ allowing comparison of the base pair and helical parameters predicted by the *ff*s with experimental X-ray crystallography values. We find that none of the tested *ff*s are able to fully reproduce the hybrid structure. A significant challenge arises from the inability of many *ff*s to sample the experimental C3′-endo pucker of the DNA strand, resulting in shifts of all major helical parameters towards more “B-form-like” geometries, even in sequences where experimental data clearly indicate dominance of the A-form. On the other hand, the simulations correctly reproduce the anisotropic buckle of the hybrids. The data presented herein offer insights into the MD simulation description of the hybrids, complement existing experimental data and outline potential opportunities for further *ff* adjustments. Furthermore, the results can guide the selection of utilizable *ff* combination for future MD studies of DNA/RNA hybrids and their complexes with proteins.

## Methods

### Selection of initial structures

We have used the X-ray structure of B-DNA dodecamer (Dickerson-Drew dodecamer (DD); PDB: 1BNA)^54^ (Figure 1) as starting structure for simulations of B-DNA duplex (DD_DNA). A-RNA (DD_RNA) and hybrid (DD_hybrid) duplexes of identical sequence were prepared with Nucleic Acid Builder.^55^ Thymines were replaced with uracils for the RNA strands and the initial helical geometry of the modeled duplexes corresponded to A-form. We also prepared hybrid duplex structure in B-form to verify the convergence of helical forms in simulations. In addition to the DD structures, simulations were also performed using the X-ray structure of the Polypurine tract hybrid duplex (PPT) (Figure 1).^13^ Lastly, to explore performance of a sequence radically different from PPT, some simulations were also performed with modeled hybrid duplex possessing 0% deoxypyrimidine content (low_dPyr), with randomized RNA strand sequence of 5′-CCUCUCUCUCCC-3′.

**Figure 1.**
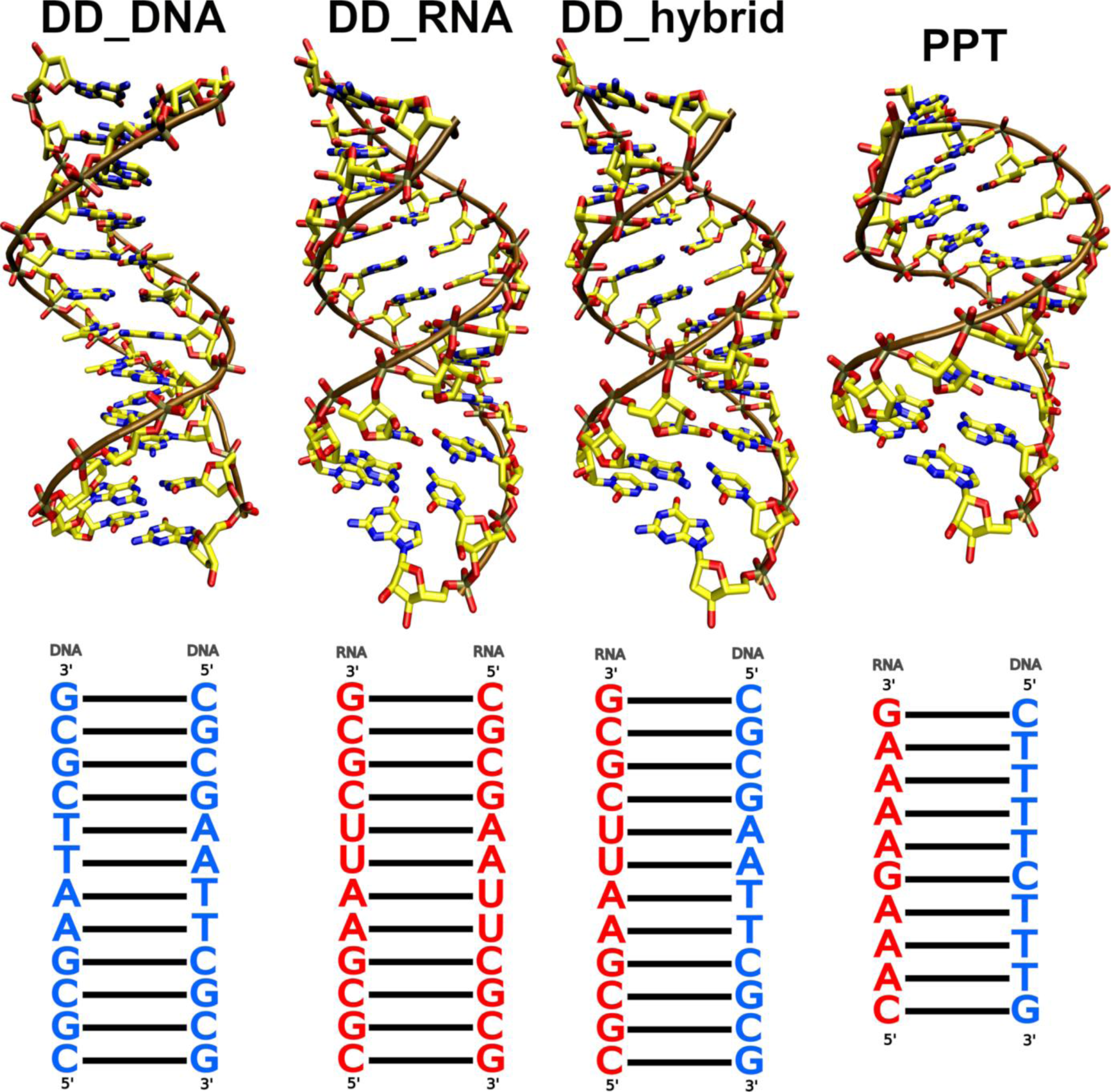
The Dickerson-Drew dodecamer (DD) and Polypurine tract hybrid duplex (PPT) structures and sequences. The DNA and RNA sequence letters are colored blue and red, respectively.

### System building and simulation protocol – the AMBER and CHARMM36 *ff*s

From the AMBER family of *ff*s, we tested the OL15,^56^ OL21,^23^ bsc1,^27^ DES-Amber,^25^ and Tumuc1^28^ *ff*s for DNA and OL3,^22^ ROC-RNA,^26^ and DES-Amber^25^ *ff*s for RNA. The tLeap program of AMBER 22^57^ was used to generate initial files for all *ff*s except DES-Amber (see below) and simulations were subsequently performed in AMBER22 using the pmemd.MPI and pmemd.cuda^58^ programs for equilibration^59^ and production simulations, respectively. The duplexes were placed in an octahedral box of SPC/E^60^ water molecules with minimal distance of 12 Å between the solute and the box border. KCl ions parametrized by Joung&Cheatham^61^ were added to neutralize the systems and to establish excess-salt ion concentration of 0.15 M. The production simulations were run for 3 μs and three independent trajectories were obtained for each system and *ff* combination. We used SHAKE^62^ and hydrogen mass repartitioning^63^ to allow 4 fs integration step. Long-range electrostatics were treated with particle mesh Ewald^64^ and periodic boundary conditions were applied. The distance cut-off for Lennard-Jones interactions was set to 9 Å. The production simulations were performed in constant pressure ensemble with pressure and temperature being regulated with Monte Carlo barostat and Langevin thermostat,^65^ respectively. In all simulations, we have applied 1 kcal/mol structure-specific HBfix (sHBfix) potential^66^ between donors and acceptors to stabilize the Watson-Crick H-bonds in the terminal base pairs, thus essentially eliminating the terminal base pair fraying. Although a minor population of frayed terminal base pairs is to be realistically expected, its estimated thermodynamic convergence in MD simulations is far beyond ten microseconds.^67^ This would lead to major sampling uncertainty and complicate the *ff* comparison. The problem has been completely circumvented by increasing stability of the terminal base pairs.

As the latest DES-Amber parameters for nucleic acids were unavailable for AMBER at the time, we have used Gromacs2018^68^ to perform all DES-Amber^25^ simulations, utilizing the parameters published in Ref.^25^ and provided by the authors on our request. The recommended TIP4P-D water model^69^ along with CHARMM22 ions^69^ was utilized in all DES-Amber simulations. Note that the Gromacs DES-Amber parameters^25^ for DNA and RNA as provided by the authors contain overlapping atomic types between the two biopolymers and therefore do not natively allow hybrid simulations. To circumvent this, we modified all the DNA atomic types and the associated parameters by adding “D” prefix to globally distinguish them from RNA-specific parameters.

We have likewise used Gromacs2018 to perform all the CHARMM36^47–48^ simulations, utilizing the CHARMM-modified TIP3P water molecules and CHARMM36 KCl parameters for ions. Due to the different molecular dynamics engine, the simulation protocol of the DES-Amber and CHARMM36 simulations somewhat differed from the ones calculated with AMBER22. Namely, the simulations utilized PLUMED 2.5^70^ to implement the sHBfix for the terminal base pairs (see above).^71^ The simulations were also performed in a rhombic dodecahedral box. Bonds involving hydrogens were constrained using the LINCS algorithm.^72^ The cut-off distance for the direct space summation of the electrostatic interactions was 12 Å and the simulations were performed at 298 K using the stochastic velocity rescale thermostat.^73^ The other settings, equilibration protocols, and system building choices were the same as in the AMBER22 simulations.

### System building and simulation protocol – DRUDE and AMOEBA polarisable *ff*s

For the simulations with CHARMM Drude19^49–51, 74^ and AMOEBA18^52–53^ polarizable *ff*s, the systems were pre-built in AMBER22, followed by minimization and equilibration using the OL21/OL3 *ff*s. In case of Drude, the resulting structures were transformed into the Drude polarizable model using the CHARMM software (version 44b1).^75^ During the conversion process, Drude particles were introduced for all heavy atoms and the lone pairs associated with each hydrogen acceptor. The SPC/E water molecules were converted into the polarizable SWM4-NDP model.^76^ Each system was then subjected to minimization and equilibration using the NAMD 2.13 package.^77^ First, we performed 1000-step minimization of the waters and ions while keeping the nucleic acid atoms restrained by a harmonic 500 kcal/mol/Å^2^ positional restraint. Keeping the duplexes restrained, this was followed by 500 ps-long MD simulation under constant pressure conditions, to relax the overall density of the systems. Afterwards, the duplexes were relaxed through several minimization cycles, gradually reducing the strength of the positional restraints placed on the nucleic acid atoms as outlined in Ref.^78^. Finally, the systems were heated under constant volume conditions for 100 ps, and then equilibrated again under constant pressure conditions for additional 100 ps. PLUMED 2.7^70^ was used to implement the sHBfix for the terminal base pairs. The obtained structures were then utilized for three separate 1-μs-long production simulations at 298 K in OpenMM 8.0.^79^ Drude Langevin integrator^80–81^ was utilized with a time step of 1 fs. The pressure was maintained at 1 bar utilizing the Monte Carlo barostat.^82^ The covalent bonds involving hydrogens were kept rigid using the SHAKE^62^ and SETTLE^83^ algorithms for solute and waters, respectively. A constraint of 0.2 Å was applied to limit the length of Drude-nuclei bonds. Electrostatic interactions were treated using the Particle-Mesh Ewald method (PME)^84^ with a 12 Å cutoff for the real space term. Non-bonded interactions were truncated at 12 Å, using a switching function from 10 to 12 Å.

The AMOEBA simulations were performed in GPU-accelerated Tinker version.^85^ The pre-built and equilibrated initial structures (see above) were converted into xyz coordinates and minimized in 10 000 steps using the steepest descent method. This was followed by heating the systems to 300 K and equilibrating the pressure to 1 bar before the production simulations were run for 1 μs in constant volume ensemble. RESPA integrator was used with the integration step of 1 and 2 fs for minimization/equilibration and production simulations, respectively. Stochastic velocity rescale thermostat^73^ and Monte Carlo barostat were used to maintain temperature and pressure, respectively. The applied real-space cutoff for electrostatics and van der Waals was 7 and 12 Å, respectively. All other settings were set to default. As Tinker did not at the time allow use of sHBfix or patching with Plumed code, we instead stabilized the H-bonds of the terminal base pairs by applying a linearly growing penalty function, with a force constant of 10 kcal/mol/Å for hydrogen-acceptor distances greater than 2.5 Å.

### Analyses

Cpptraj^86^ and VMD^87^ were used to analyse and visualize the trajectories. Molecular figures and graphs were prepared with Povray and gnuplot, respectively. The helical parameters were calculated using the *nastruct* command in cpptraj, which utilizes the same algorithms and definitions of helical parameters and groove sizes as the X3DNA.^88–90^ We used the Altona et al. definition to calculate the sugar pucker.^91^ The terminal base pairs as well as the base pair steps between the terminal and subterminal base pairs were not included in the analyses. For the DD structures, we evaluated the base pair parameters as separate averages of the AT(U) and GC base pairs. For the other sequences, averages of all non-terminal base pairs were calculated. For the CHARMM36 simulations, simulation frames with disrupted non-terminal base pairs were excluded from the analyses (see below). The histogram analyses were performed in cpptraj with bin sizes ranging from 0.1 to 5 depending on the range of absolute values observed for the individual parameters (see Supporting Information). All histograms were normalized by cpptraj so that the sum of the bins equals one and the presented graphs are shown with cubic spline smoothing applied by Gnuplot. Unless specified otherwise, the presented values correspond to combined simulation ensembles of the three simulations performed for each system and *ff* combination. The first 500 ns of each production simulation were considered an extended equilibration period and were not included in the analyses. All the trajectories were visually inspected.

## Results and Discussion

Below we present an extensive set of MD simulations involving A-RNA, B-DNA, and hybrid DNA/RNA duplexes (120 independent simulations in total with a cumulative time of 312 μs, see Table 1). Average simulation values of the helical parameters relevant for the study are given in Table 2 and Supporting Information Table S2. To summarize, by simulating A-RNA, B-DNA and hybrid helices of the same sequence (Figure 1 and Table 1), we were able to capture the changes in helical form balance among the three duplexes (Table 2). The details differed significantly depending on the utilized *ff*s, but in general the individual helical parameters of the hybrid duplexes were either closer but not identical to the A-form values (inclination, helical twist, propeller twist, minor groove width), near midway between values characteristic for A- and B-forms (x-displacement, roll, slide, helical rise), or they possessed unique values divergent from both forms (buckle, tilt, tip) (Table 2 and Supporting Information Table S2). By comparing our simulations with the helical values observed experimentally for the PPT, we then evaluated how accurately the individual *ff* combinations reproduce this hybrid structure (Table 2 and Supporting Information Table S2). The following text details the key observations derived from our analyses.

**Table 1.** List of simulations.

| duplex | force field<br>DNA \| RNA |  | water model | simulations ×<br>length [μs] |
| --- | --- | --- | --- | --- |
| non-polarizable force fields |  |  |  |  |
| DD_DNA | OL21 | - | SPC/E | 3 × 3 |
| DD_DNA | OL15 | - | SPC/E | 3 × 3 |
| DD_DNA | bsc1 | - | SPC/E | 3 × 3 |
| DD_DNA | Tumuc1 | - | SPC/E | 3 × 3 |
| DD_DNA | DES-Amber | - | TIP4P-D | 3 × 3 |
| DD_DNA | CHARMM36 | - | TIP3P | 3 × 3 |
| DD_RNA | - | OL3 | SPC/E | 3 × 3 |
| DD_RNA | - | ROC-RNA | SPC/E | 3 × 3 |
| DD_RNA | - | DES-Amber | TIP4P-D | 3 × 3 |
| DD_RNA | - | CHARMM36 | TIP3P | 3 × 3 |
| DD_hybrid | OL21 | OL3 | SPC/E | 3 × 3 |
| DD_hybrid<sup>a</sup> | OL21 | OL3 | SPC/E | 3 × 3 |
| DD_hybrid | OL21 | OL3 | OPC | 3 × 3 |
| DD_hybrid | OL15 | OL3 | SPC/E | 3 × 3 |
| DD_hybrid<sup>b</sup> | OL15 | OL3 | SPC/E | 3 × 3 |
| DD_hybrid<sup>c</sup> | OL15 | OL3 | SPC/E | 3 × 3 |
| DD_hybrid | bsc1 | OL3 | SPC/E | 3 × 3 |
| DD_hybrid | bsc1 | OL3 | OPC | 3 × 3 |
| DD_hybrid | OL21 | ROC-RNA | SPC/E | 3 × 3 |
| DD_hybrid | Tumuc1 | OL3 | SPC/E | 3 × 3 |
| DD_hybrid | Tumuc1 | ROC-RNA | SPC/E | 3 × 3 |
| DD_hybrid | DES-Amber | DES-Amber | TIP4P-D | 3 × 3 |
| DD_hybrid | CHARMM36 | CHARMM36 | TIP3P | 3 × 3 |
| DD_hybrid<sup>d</sup> | CHARMM36 | CHARMM36 | TIP3P | 3 × 3 |
| PPT | OL21 | OL3 | SPC/E | 3 × 3 |
| PPT | OL15 | OL3 | SPC/E | 3 × 3 |
| PPT | bsc1 | OL3 | SPC/E | 3 × 3 |
| PPT | OL21 | ROC-RNA | SPC/E | 3 × 3 |
| PPT | Tumuc1 | OL3 | SPC/E | 3 × 3 |
| PPT | DES-Amber | DES-Amber | TIP4P-D | 3 × 3 |
| PPT | CHARMM36 | CHARMM36 | TIP3P | 3 × 3 |
| low_dPyr | OL21 | OL3 | SPC/E | 3 × 3 |
| polarizable force fields |  |  |  |  |
| DD_DNA | DRUDE | - | SWM4-NDP | 3 × 1 |
| DD_DNA | AMOEBA | - | AMOEBA | $3 \times 1$ |
| DD_RNA | - | DRUDE | SWM4-NDP | $3 \times 1$ |
| DD_RNA | - | AMOEBA | AMOEBA | $3 \times 1$ |
| DD_hybrid | DRUDE | DRUDE | SWM4-NDP | $3 \times 1$ |
| DD_hybrid | AMOEBA | AMOEBA | AMOEBA | $3 \times 1$ |
| PPT | DRUDE | DRUDE | SWM4-NDP | $3 \times 1$ |
| PPT | AMOEBA | AMOEBA | AMOEBA | $3 \times 1$ |

<sup>a</sup>The DNA nucleotide puckers were restrained to C3'-endo region (pseudorotation values of $-10^\circ$ to $40^\circ$ ) by five flat-well dihedral restraint potential functions applied on each 2-deoxyribose (see Supporting Information).
<sup>b</sup>The initial structure of the hybrid corresponded to the B-form helix (see Methods).
<sup>c</sup>The $\chi$ dihedral potentials of the OL15 DNA *ff* were replaced with the RNA $\chi$ dihedral potentials from OL3 *ff*. See Ref.<sup>92</sup> for more details.
<sup>d</sup>2 kcal/mol stabilizing sHBfix potential was applied on every base pairing H-bond.

**Table 2.** Selected helical parameters observed in the MD simulations.

| duplex | force field<br>DNA / RNA | buckle<br>AT / GC | propeller<br>AT / GC | inclination | x-displacement | roll |
| --- | --- | --- | --- | --- | --- | --- |
| **Non-polarizable force fields** |  |  |  |  |  |  |
| DD_DNA | OL21 / - | -0.1/0.0 | -17.5/-5.9 | 2.7 | -0.3 | 1.3 |
| DD_DNA | OL15 / - | -0.1/0.0 | -17.6/-5.9 | 3.0 | -0.4 | 1.5 |
| DD_DNA | bsc1 / - | 0.3/-0.1 | -15.8/-4.2 | 2.7 | -0.8 | 1.1 |
| DD_DNA | Tumuc1 / - | 0.0/0.0 | -13.5/-5.9 | -0.3 | 0.0 | -0.5 |
| DD_DNA | DES-Amber / - | 0.0/0.0 | -15.7/-7.9 | 5.9 | -1.6 | 3.4 |
| DD_DNA | CHARMM36 / - | 0.0/0.0 | -18.5/-7.4 | 10.0 | -0.7 | 5.6 |
| DD_RNA | - / OL3 | 0.1/-0.1 | -14.2/-12.4 | 17.0 | -4.3 | 9.5 |
| DD_RNA | - / ROC-RNA | -0.0/-0.1 | -13.5/-10.4 | 12.4 | -4.3 | 6.8 |
| DD_RNA | - / DES-Amber | -0.0/0.0 | -11.9/-8.9 | 12.2 | -5.2 | 6.3 |
| DD_RNA | - / CHARMM36 | 1.1/-0.8 | -14.8/-14.7 | 18.4 | -4.5 | 10.3 |
| DD_hybrid | OL21 / OL3 | -3.9/-9.2 | -15.4/-10.8 | 12.5 | -2.9 | 7.2 |
| DD_hybrid<sup>a</sup> | OL21 / OL3 | -2.7/-1.9 | -16.1/-14.0 | 16.8 | -3.9 | 9.4 |
| DD_hybrid | OL21 / OL3<br>(OPC water) | -4.0/-9.7 | -14.9/-10.5 | 12.2 | -3.1 | 6.9 |
| DD_hybrid | OL15 / OL3 | -4.2/-9.0 | -15.4/-10.7 | 12.6 | -2.9 | 7.1 |
| DD_hybrid<sup>b</sup> | OL15 / OL3 | -3.3/-8.1 | -15.7/-10.5 | 12.3 | -2.8 | 7.0 |
| DD_hybrid<sup>c</sup> | OL15 / OL3 | -1.9/-7.2 | -13.6/-10.2 | 12.2 | -3.3 | 6.8 |
| DD_hybrid | bsc1 / OL3 | -2.4/-7.7 | -14.0/-10.0 | 12.1 | -3.1 | 6.8 |
| DD_hybrid | bsc1 / OL3<br>(OPC water) | -2.6/-8.2 | -13.5/-9.6 | 11.5 | -3.2 | 6.4 |
| DD_hybrid | OL21 / ROC-RNA | -8.3/-9.7 | -15.6/-10.1 | 10.1 | -2.6 | 5.6 |
| DD_hybrid | Tumuc1 / OL3 | -6.6/-12.3 | -9.9/-8.9 | 11.0 | -3.2 | 6.1 |
| DD_hybrid | Tumuc1 / ROC-RNA | -7.1/-11.4 | -11.5/-8.6 | 8.6 | -2.7 | 4.8 |
| DD_hybrid | DES-Amber / DES-Amber | -5.6/-8.0 | -12.6/-9.1 | 11.4 | -4.0 | 6.1 |
| DD_hybrid | CHARMM36 / CHARMM36 | -6.6/-9.7 | -14.0/-13.0 | 16.1 | -3.5 | 9.2 |
| DD_hybrid<sup>d</sup> | CHARMM36 / CHARMM36 | -6.1 / -11.0 | -13.7 / -13.2 | 17.0 | -3.6 | 9.6 |
| PPT | *X-ray structure*<sup>e</sup> | -4.6 | -10.3 | 14.3 | -3.9 | 8.4 |
| PPT | OL21 / OL3 | -3.7 | -14.4 | 6.3 | -2.5 | 3.5 |
| PPT | OL15 / OL3 | -5.0 | -13.7 | 7.2 | -2.8 | 4.0 |
| PPT | bsc1 / OL3 | -4.0 | -12.1 | 6.3 | -2.9 | 3.5 |
| PPT | OL21 / ROC-RNA | 0.4 | -17.7 | 4.4 | -1.8 | 2.4 |
| PPT | Tumuc1 / OL3 | -2.4 | -9.0 | 2.2 | -2.6 | 1.2 |
| PPT | DES-Amber / DES-Amber | -8.3 | -12.4 | 8.3 | -3.4 | 4.4 |
| PPT | CHARMM36 / CHARMM36 | -6.4 | -11.4 | 13.7 | -4.1 | 7.4 |
| low_dPyr | OL21 / OL3 | -3.5 | -11.5 | 15.3 | -3.4 | 8.3 |
| **Polarizable force fields** |  |  |  |  |  |  |
| DD_DNA | DRUDE / - | 0.0/-0.1 | -11.7/-5.0 | 3.1 | -0.8 | 1.4 |
| DD_DNA | AMOEBA / - | -0.0/0.2 | -14.2/-2.7 | -0.2 | -0.4 | -0.2 |
| DD_RNA | - / DRUDE | 0.0/0.0 | -20.8/-14.2 | 11.9 | -3.2 | 7.0 |
| DD_RNA | - / AMOEBA | 0.2/-0.1 | -5.1/-4.7 | 8.6 | -5.1 | 4.7 |
| DD_hybrid | DRUDE / DRUDE | -4.0/-6.6 | -14.2/-9.1 | 8.0 | -2.3 | 4.8 |
| DD_hybrid | AMOEBA / AMOEBA | -4.9/-5.7 | -4.4/-4.0 | 9.1 | -4.9 | 4.8 |
| PPT | DRUDE / DRUDE | 1.5 | -10.6 | 0.2 | -1.9 | 0.1 |
| PPT | AMOEBA / AMOEBA | -6.2 | -7.1 | 5.3 | -3.4 | 2.8 |

<sup>a</sup>The DNA nucleotide puckers were restrained to C3'-endo region (pseudorotation values of -10° to 40°) by five flat-well dihedral restraint potential functions applied on each 2-deoxyribose (see Supporting Information).
<sup>b</sup>The initial structure of the hybrid corresponded to the B-form helix (see Methods).
<sup>c</sup>The $\chi$ dihedral potentials of the OL15 DNA *ff* were replaced with the RNA $\chi$ dihedral potentials from OL3 *ff*. See Ref.<sup>92</sup> for more details.
<sup>d</sup>2 kcal/mol stabilizing sHBfix potential was applied on every base pairing H-bond.
<sup>e</sup>Average values observed in the PPT experimental structure.<sup>13</sup> See Supporting Information Table S1 for values observed for other experimental structures of the hybrids.

### The AMBER family of *ff*s did not reproduce the C3′-endo pucker of the DNA strand of the hybrid duplexes

In all hybrid simulations using the non-polarizable AMBER *ff*s, we observed the DNA nucleotides possessing exclusively C2′-endo puckers (∼135–180°) (Figure 2) along with N-glycosidic χ dihedrals in the high-*anti* region (∼250°), both characteristic features of the B-form duplexes. This simulation result stands in stark contrast to experiments which instead indicate C3′-endo pucker (∼-10–40°) and χ *anti* (∼200°); exclusively in case of X-ray structures and as a population mixture of C2′/C3′-endo puckers in NMR data. The simulations showed slightly lower pucker values for the DNA in hybrids than in pure DNA, but still well within the C2′-endo region. We suggest these observations are related to the known difficulties with addressing the A-DNA/B-DNA balance and transitions in MD simulations using the AMBER *ff*s, where the absence of C3′-endo pucker for DNA is also problematic.^92–93^

**Figure 2.**
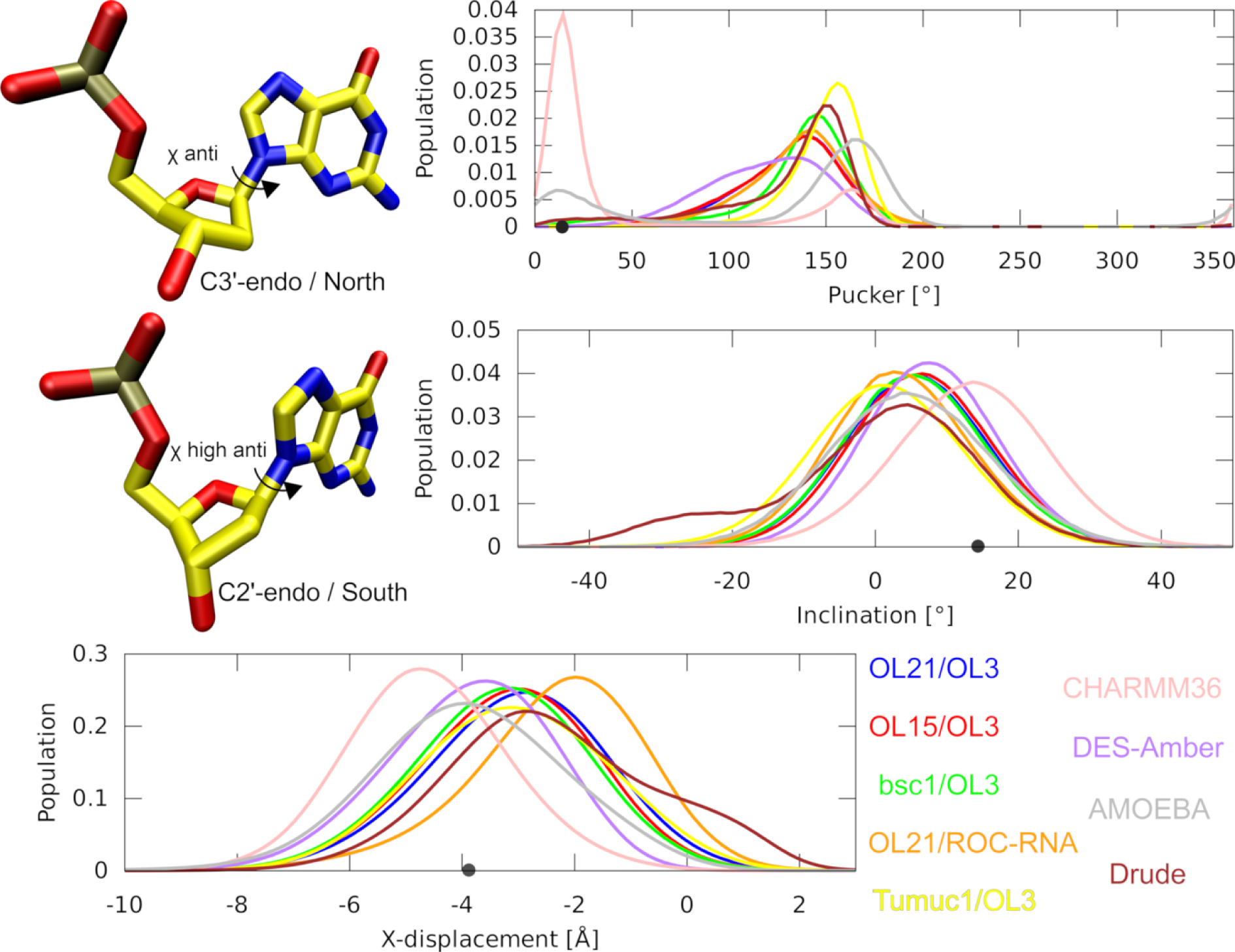
Histograms of the DNA sugar puckers, inclination and x-displacement in MD simulations of the PPT structure. Example of a nucleotide possessing C3′-endo and C2′-endo puckers, respectively, is shown at top left corner. The individual *ff* combinations are color-coded in the graphs according to the legend in the bottom right corner. Black dots on the x-axes indicate average experimental values.

According to the solution NMR measurements, the sugar pucker of the DNA nucleotides in hybrids exists in a state of C2′/C3′-endo dynamic equilibrium.^14–15, 94^ Therefore, we conclude the complete inaccessibility of the C3′-endo pucker for the DNA strand in our hybrid simulations using all the AMBER *ff*s points to a significant imbalance. The absolute preference for C2′-endo leads to majority of the helical parameters shifting closer to the B-form (Table 2 and Supporting Information Table S2). Most significant is the reduction of inclination and roll, which in case of the PPT structure clearly contradicts the experimental data (Table 2). The inclination and roll are interrelated parameters in global helical and local base pair steps coordinate systems, respectively.^95–96^ We also noticed that the backbone dihedral angles of the hybrids correspond closely to the pure duplexes. In other words, the dihedrals observed for the hybrid RNA strand were virtually identical to the A-RNA duplex while the hybrid DNA strand was nearly identical to the B-DNA duplex, showing no discernible adaptation to the unique structural environment of the hybrids (Figure 3).

**Figure 3.**
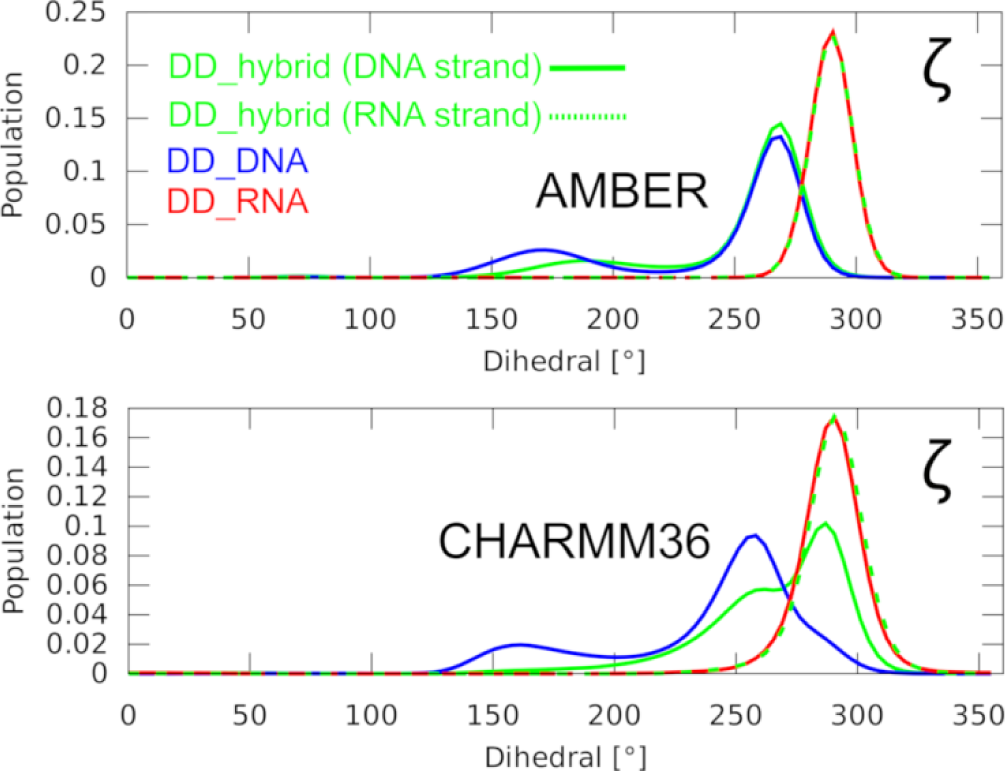
Histograms of the dihedral angles in MD simulations of the DD structures. Distribution of the ζ (zeta) dihedral angle in pure DNA and RNA and the hybrid DD structures, using the OL21/OL3 (top) and CHARMM36 *ff*s (bottom). The visualized datasets are representative of the development observed also for the other dihedrals. The top graph is representative of a performance seen also with the other AMBER *ff*s.

The C3’-endo pucker of the hybrid DNA strand can be safely enforced by restraints (Table 1 and Supporting Information Figure S1) without undesired side-effects on the duplex stability. It was tested for the OL21/OL3 *ff* combination where it leads to significant shifts of the helical parameters towards the A-form (Table 2). We suggest that restraining the DNA sugar pucker in hybrids to C3′-endo might be helpful in simulations of hybrids using the AMBER *ff*s when such pucker is required, e.g. for correct interaction geometry with a protein or when detailing some catalytic mechanisms. In contrast, replacing the DNA χ dihedral potentials in OL15 with the RNA potentials from OL3 *ff*, as recently tested in Ref.^92^ to improve the A/B-form balance of the DNA simulations, provided negligible improvements for the hybrid (Supporting Information Figure S1). We also note the absence of C3′-endo DNA pucker in AMBER simulations might not always be detrimental for the helical parameters as we observed relatively high inclination in the low_dPyr simulations (see Methods and Table 2), despite complete lack of the C3′-endo population on the DNA strand (Supporting Information Figure S1).

### CHARMM36 *ff* reproduces the C3′-endo pucker of the hybrids but has trouble to keep stable base pairing

The simulations using the CHARMM36 *ff* accurately reproduced the experimental PPT hybrid structure geometry, including the C3′-endo pucker of the DNA strand (Figure 2) and the inclination (Table 2). Contrasting the AMBER *ff*s, we also observed the DNA strand of the hybrid mostly possessing dihedrals typical for pure A-RNA (Figure 3), rationalizing the shift of the helical parameters toward the A-form (Table 2). The CHARMM36 performance we observed for the hybrids is very good and in agreement with the previous studies utilizing the CHARMM *ff*.^35–36, 97^ However, we also noticed large instabilities of the base pairing in all our CHARMM36 simulations (Supporting Information Figure S3). Namely, several consecutive base pairs near the terminus were temporarily or permanently disrupted in CHARMM36 simulations (both hybrids and pure duplexes). These large instabilities occurred despite the terminal base pairs being supported with the sHBfix (see Methods), which was sufficient to almost entirely prevent terminal base pair fraying in simulations using the AMBER *ff*s. While some degree of fraying is to be expected,^67^ we suggest such large scale instability on a microsecond timescale (up to 30% of simulation frames were affected by this), including the thermodynamically stable GC/CG base pairs (Supporting Information Figure S3), likely reflects imbalance within the CHARMM36 *ff* rather than genuine biomolecular dynamics.

A possible way to overcome this problem and still benefit from the satisfactory performance of CHARMM36 *ff* for hybrids is to apply the sHBfix potentials^66^ to increase the stability of every base pair and not just the terminal ones (Table 1). Our CHARMM36 simulations where we have applied sHBfix in such fashion revealed negligible instability of the base pairs while fully maintaining the quite accurate description of the hybrid structure (Table 2). The advantage of stabilizing the base pairing with sHBfix instead of standard distance restraints is that temporary disruptions of the base pairing and breathing motions are still allowed to happen, albeit with lower frequency, leading to a more natural dynamics.^98^

### The polarizable *ff*s can populate C3′-endo pucker for DNA nucleotides but significantly underestimate the inclination

The polarizable DRUDE and AMOEBA *ff*s sample a minor population of the C3′-endo pucker in the DNA strand of the hybrid systems (Figure 2). Notably, in the DD_hybrid simulations using the AMOEBA *ff*, the C3′-endo population was even dominant (Supporting Information Figure S1). However, surprisingly, the population of the DNA C3′-endo pucker did not translate into increased base pair inclination of the hybrids (Table 2). In fact, the inclination was generally lower than with the AMBER family of *ff*s where the C3′-endo pucker for DNA was completely absent and where higher inclination could be achieved by restraining the C3′-endo pucker (see above). The other helical parameters were also farther away from the experimental values in comparison with the simulations using the non-polarizable *ff*s. We noticed the polarizable and non-polarizable *ff*s populating different backbone dihedral angles, which might rationalize the suboptimal reproduction of the helical parameters despite the C3′-endo pucker being accessible for the DNA strand (Supporting Information Figure S4).

### Additional comments on specific force fields

We have also observed some problems with specific *ff*s and their combinations (Table 1). Namely, the ROC-RNA *ff* reversibly but significantly populated the α(trans)/γ(trans) dihedral region of the RNA backbone (Figure 4a), both in pure RNA and hybrids. Such large population of this dihedral angle combination was not seen for any other *ff.* It might have contributed to the low inclination and drift towards the B-form observed for the hybrids with this *ff* (Table 2), as by excluding the frames with α(trans)/γ(trans) states from the ensemble, the inclination of the PPT nearly doubled. Secondly, while the Tumuc1 *ff* performed well for the B-DNA simulations, when combined with either OL3 or ROC-RNA we observed a visible drift towards B-form for the hybrids, once again reducing both the inclination and x-displacement below their experimental values (Table 2 and Figure 2). Lastly, the DES-Amber *ff* revealed a complete lack of backbone BII-states in pure B-DNA, as admitted also in the original paper.^25^ Consequently, the BII-states were absent also for the DNA strand in the hybrids (Figure 4b). The DES-Amber *ff* also revealed an unusual population of non-native pucker values in C4′-exo/O4′-endo region for DNA nucleotides in both B-DNA and hybrids (Figure 4c). This may have shifted some of the hybrid helical parameters closer to the experimental values as the pucker value was lowered closer to the experimentally indicated C3′-endo state, but did not reach it (Figure 4c and Table 2). In any case, C4′-exo/O4′-endo puckers are non-native for B-DNA and their presence in simulations is likely undesirable for both B-DNA and the hybrids.

**Figure 4.**
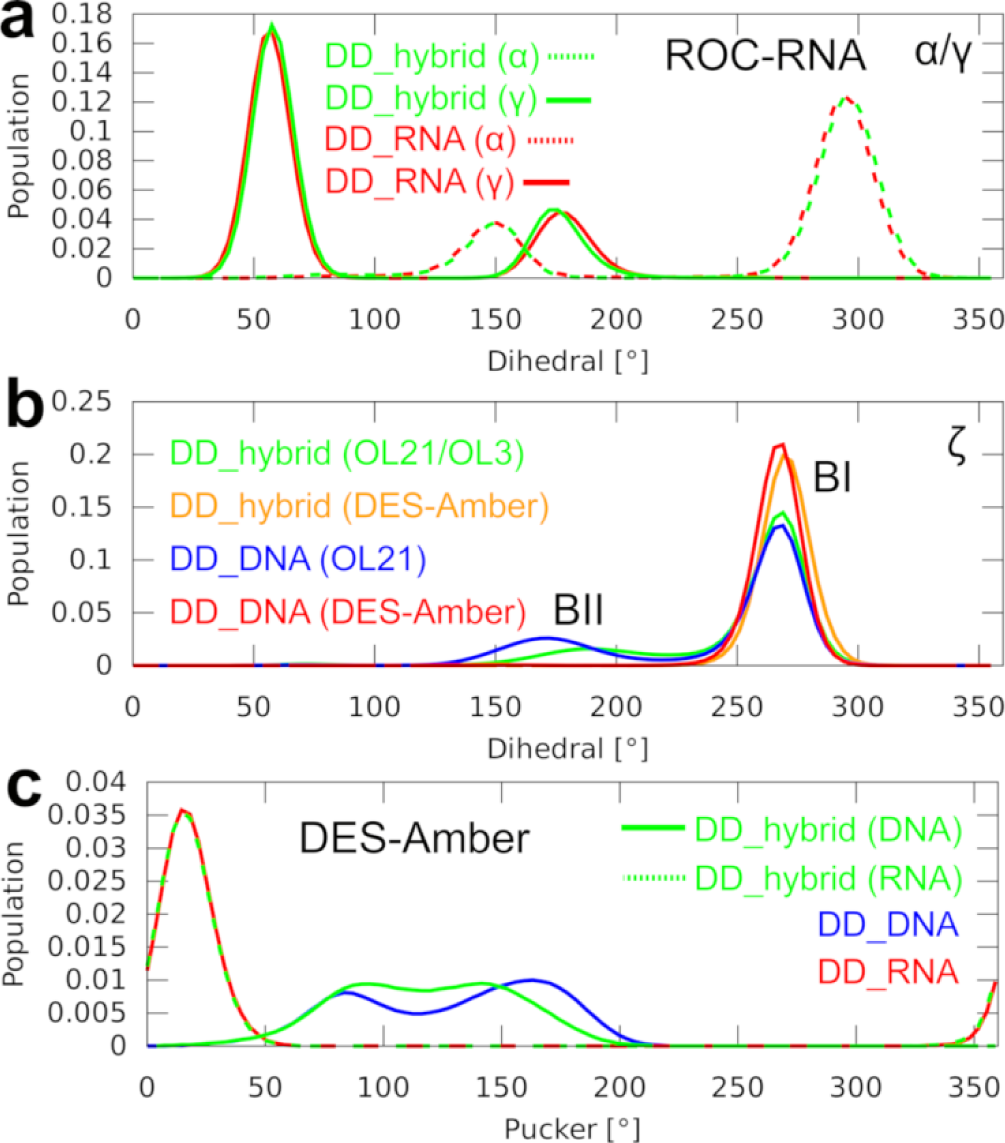
Histograms of the sugar puckers and dihedral angles in MD simulations of the DD structures. **a)** α and γ dihedrals observed for the RNA strand in simulations of the DD_RNA and DD_hybrid structures using the ROC-RNA *ff* (combined with OL21 *ff* for the hybrid). **b)** ζ dihedral angle in DD_DNA and DD_hybrid structures, using the OL21/OL3 and DES-Amber *ff*s. Correlated behavior was observed for the ε dihedral as it pertains to the BI/BII B-form substrates. **c)** Sugar puckers observed for the DD structures using the DES-Amber *ff*.

In general, we also recommend careful verification that the hybrid systems are correctly built as some *ff*s (e.g. DES-Amber *ff*, see Methods) can contain overlapping atomic types for the DNA and RNA nucleotides. A good practice is to always check that the same potential energies of the individual strands are obtained for systems built with lone DNA and RNA strands and a single set of parameters loaded, and when the same molecules are part of the hybrid. Disagreement between the calculated energies indicates an overlap between the parameters.

### Water model seems to have minimal influence

We have utilized water models recommended by the *ff* developers, provided such recommendations were given in the original papers. In other cases, we have used the SPC/E water model,^60^ which is well tested for simulations of nucleic acids in general. With certain *ff* combinations, we have also tested the OPC water model^99^ for the hybrid simulations (see Methods and Table 1). The results of these simulations indicated only a minor sensitivity to the water model used, with the solute *ff* proving to be the decisive factor. Specifically, the hybrid simulations using the OPC water revealed a minor but systematic drop in inclination compared to the SPC/E simulations (Table 2 and Supporting Information Figure S2). While this further exacerbates the undesirable reduction of inclination we observe for the hybrids, it is a very minor effect. Although we did not test the water models systematically, we tentatively suggest that for simulations of hybrid duplexes all the water models commonly employed for simulations of nucleic acids (e.g. TIP3P, SPC/E and OPC) are reasonable choices. The results are only moderately sensitive to the water model choice, echoing similar conclusions made for simulations of A-RNA duplexes.^78, 96, 100^ Therefore, it might be best to follow the recommendation made by the authors of the particular nucleic acid *ff*, provided any are given. Obviously, this can lead to incompatible water model requirements for some DNA and RNA *ff* combinations. For instance, the DES-Amber *ff* utilizes the TIP4P-D water model along with the CHARMM22 ions, which is however an untested setting with the other AMBER *ff*s.

### Hybrid duplexes display buckle anisotropy

An intriguing difference between the pure duplexes and hybrids was observed for the *buckle*, which describes the V-shaped deformation of the base pairs (Figure 5a). Namely, the distribution of the buckle peaked around 0° for both A-RNA and B-DNA, meaning that on average the base pairs did not have a preferred direction for the V-shaped deformation as they fluctuated in both directions equally. In contrast, the buckle peaked at ∼ −5° for the hybrids, making the base pairs buckled in the 3′ direction of the RNA strand on average (Table 2 and Figure 5). Examination of the helical parameters in nine different hybrid duplexes deposited in the PDB database revealed the buckle anisotropy to be a universal structural rule for the hybrids (Supporting Information Table S1). All tested *ff*s managed to qualitatively reproduce this feature. Still, the buckle value significantly varied among the tested *ff* combinations (Table 2). While seemingly a genuine structural effect, we suspect that some of the *ff*s could be exaggerating it in order to relieve potential conflicts between the RNA and DNA *ff* parameters. This was most visible for the Tumuc1/OL3, OL21/ROC-RNA, and DES-Amber *ff* combinations (Figure 5b). Lastly, note that the tilt and tip parameters were similarly shifted for the hybrids as the buckle (Supporting Information Table S2).

**Figure 5.**
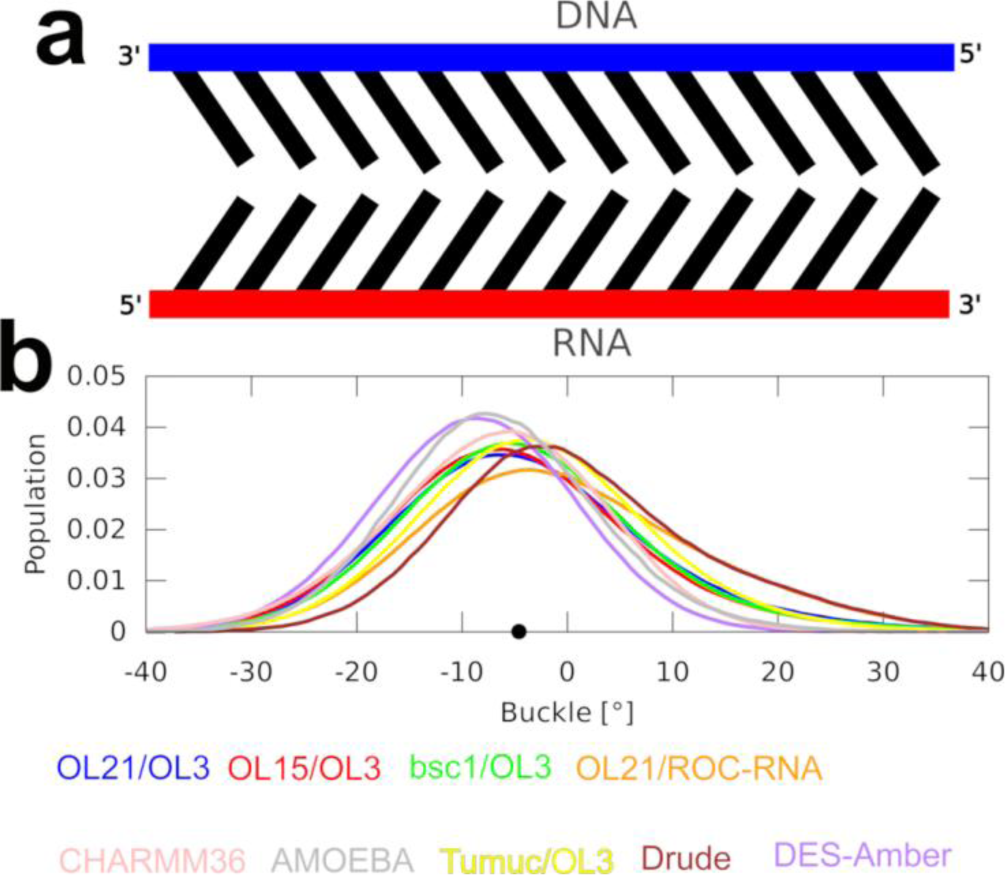
Anisotropic base pair buckle observed in the hybrid DNA/RNA experimental structures and simulations. **a)** 2D scheme of the anisotropic buckle of the hybrids. Note that the extent of the V-shaped deformation of the base pairs is greatly exaggerated for illustrative purposes. **b**) Distribution of the buckle in MD simulations of the PPT structure. Black dot on the x-axis indicates the average experimental value.

### Convergence of the helical forms

The standard timescale for our MD simulations was 3 μs (1 μs for the polarizable *ff*s) and we have executed at least three independent simulations for each system and *ff* combination. First 500 ns of each simulation were carefully monitored but not considered for the analyses described above (see Methods). Comparison of the parallel simulations revealed average RMSD between the ensembles of less than 0.1 Å, suggesting excellent convergence. In fact, we universally observed the systems approaching convergence of their helical parameters in ∼200-300 ns, with B-DNA and hybrids taking longer than the A-RNA due to sampling of the BI/BII populations in the DNA strand(s). With some *ff*s, these were sampled also in the DNA strand of the hybrid, but compared to the pure DNA, the BII dihedrals were less populated (Figure 4b). Convergence of the hybrid’s helical form was of special concern as in principle it can be initially modeled as A-form (utilized in most of our simulations; see Methods) or B-form, with the initial structure potentially affecting the results. However, by comparing the average structures obtained from simulations started from the two helical forms, we observed the RMSD of the second halves of the simulation ensembles of ∼0.1 Å. The resulting helical parameters were also very similar (Table 2), showing differences on par with the negligible differences observed among parallel simulations started from the same helical form. This confirms that the helical form of the starting structure is not affecting the results. We have likewise monitored the time development of all the individual helical parameters, dihedrals and sugar puckers described in the text, observing extensive and frequent fluctuations throughout the simulations with all *ff*s. In conclusion, we suggest that our simulations were long enough to capture all the qualitative trends associated with the individual systems and *ff*s, as well as to achieve a very good quantitative convergence.

## Conclusions

In this study, we present a comprehensive MD simulation benchmark focusing on hybrid DNA/RNA duplexes, utilizing modern, state-of-the-art nucleic acid *ff*s. Our findings suggest that none of the tested *ff*s provide a fully satisfactory performance. The OL-family of AMBER *ff*s (OL21, OL15 for DNA, and OL3 for RNA), the bsc1 (paired with OL3 for RNA) or CHARMM36, currently represent the most promising albeit far from perfect choices for simulating the hybrids. The universal limitation of all tested AMBER *ff*s was the inability to reproduce C3′-endo pucker of the DNA strand in the hybrids, accompanied by reduced helical inclination. This appears to be primarily the issue of the DNA *ff* parameters. We show that when utilizing the AMBER *ff*s for hybrid simulations, it is straightforwardly possible to restrain the C3′-endo pucker for the DNA nucleotides, which greatly improves the inclination. The CHARMM36 *ff* provided satisfactory sugar pucker distributions and inclinations, but the *ff* severely underestimates stability of the base pairs in both hybrids and pure duplexes. It leads to large populations of structures with several disrupted base pairs, not only the terminal ones. We demonstrate that this issue can be resolved by reinforcing stability of the base pairs using sHBfix, which significantly improved simulation performance with CHARMM36. With the help of sHBfix, CHARMM36 provides the best results for the hybrid duplexes. The latest versions of polarizable DRUDE and AMOEBA *ff*s demonstrated a promising ability to populate the C3′-endo pucker as a minor state and to describe the pucker transitions. However, their inclination was even lower than with the non-polarizable *ff*s, possibly requiring further *ff* refinement, perhaps of the backbone dihedral potentials. In summary, the results show that accurate simulations of hybrid DNA/RNA duplexes are challenging for contemporary *ff*s. It is apparent that dedicated parametrization efforts would likely be necessary to develop a perfect *ff* for the hybrid duplexes. At the very least, DNA/RNA hybrids represent a straightforward test system that should be considered in future studies aimed at *ff* refinement, especially of the DNA parameters. Still, satisfactory performance for the hybrid simulations can be achieved with current generation of *ff*s, provided the authors are cognizant of their specific caveats and apply the appropriate workarounds, as outlined in this work.

## Supplementary Information

Helical parameters of other X-ray and NMR structures of hybrid DNA/RNA helices, not utilized in this study. Additional helical parameters calculated for the simulated structures. Additional histograms and structural figures.

