## Supporting Information for "Comprehensive Assessment of Force-Field Performance in Molecular Dynamics Simulations of DNA/RNA Hybrid Duplexes"

### Supporting Information Text

**Size of the bins for the histograms.** The size of the histogram bins for the individual parameters was 5 (dihedrals) or 1 (inclination, buckle and x-displacement). The binning range was adjusted for each parameter so that all the populated values in every simulation could be binned.

**Flat-well distance restraint functions applied on DNA sugars to enforce C3′-endo puckers.** The following restraint functions (AMBER format) were applied on each sugar ring to maintain the sugar pucker in the range of -10 to 40°. Example for the first nucleotide of the DNA strand in DD_hybrid system is shown. Note that in principle the strength of the restraints can be adjusted and fine-tuned to obtain the desired C3′-endo/C2′-endo balance.

```
# 13 DC NU0:  (13 DC5 C4')-(13 DC5 O4')-(13 DC5 C1')-(13 DC5 C2') -15.0  15.0
 &rst    iat =   391,   393,   394,   410,
            r1 = -16.0, r2 = -15.0, r3 =  15.0, r4 =  16.0,
            rk2 =   2000.0, rk3 =   2000.0,                              /

# 13 DC NU1:  (13 DC5 O4')-(13 DC5 C1')-(13 DC5 C2')-(13 DC5 C3') -36.5  -6.5
 &rst    iat =   393,   394,   410,   408,
            r1 = -37.5, r2 = -36.5, r3 =  -6.5, r4 =  -5.5,
          /

# 13 DC NU2:  (13 DC5 C1')-(13 DC5 C2')-(13 DC5 C3')-(13 DC5 C4')  19.8  49.8
 &rst    iat =   394,   410,   408,   391,
            r1 =  18.8, r2 =  19.8, r3 =  49.8, r4 =  50.8,
```

```
/

# 13 DC NU3:  (13 DC5 C2')-(13 DC5 C3')-(13 DC5 C4')-(13 DC5 O4') -49.8 -19.8
 &rst    iat =   410,   408,   391,   393,
          r1 = -50.8, r2 = -49.8, r3 = -19.8, r4 = -18.8,
         /

# 13 DC NU4:  (13 DC5 C3')-(13 DC5 C4')-(13 DC5 O4')-(13 DC5 C1')   6.5  36.5
 &rst    iat =   408,   391,   393,   394,
          r1 =   5.5, r2 =   6.5, r3 =  36.5, r4 =  37.5,
```

### Supporting Information Tables

Table S1. Selected helical parameters observed in the hybrid duplex structures deposited in the PDB database.[a]

| Hybrid Duplexes | method | Sequence[b] | Buckle | Propeller | Inclination | x-displacement |
|---|---|---|---|---|---|---|
| 479D | X-ray | DNA/RNA | 5.70 | -11.55 | 14.03 | -3.98 |
| 404D | X-ray | RNA/DNA | -7.96 | -7.50 | 10.36 | -4.56 |
| 3SSF | X-ray | RNA/DNA | -5.10 | -16.65 | 16.36 | -3.30 |
| 1FIX | X-ray | RNA/DNA | -6.20 | -13.67 | 15.19 | -4.01 |
| 1NXR | NMR | DNA/RNA | 5.74 | -12.82 | 8.13 | -4.25 |
| 8DRH | NMR | DNA/RNA | 6.42 | -15.79 | 16.19 | -3.54 |
| 1HG9 | NMR | DNA/RNA | 2.53 | -17.16 | 17.69 | -3.55 |
| 124D | NMR | DNA/RNA | 7.75 | -12.43 | 11.54 | -3.71 |
| 1EFS | NMR | DNA/RNA | 6.32 | -6.83 | 6.48 | -3.47 |

[a]For solution NMR, the average values of the entire ensemble are given.
[b]Indicates whether DNA or RNA strand is positioned first in the PDB file. It affects the sign of the buckle value.

Table S2. Additional helical parameters observed in the MD simulations.

| Duplex | force field DNA / RNA | Minor groove width AT / GC | Slide | Tilt | Helical rise | Tip | Helical twist |
|---|---|---|---|---|---|---|---|
| **Non-polarizable force fields** | | | | | | | |
| DD_DNA | OL21 / - | 10.35/11.57 | 0.02 | 0.00 | 3.27 | 0.00 | 35.93 |
| DD_DNA | OL15 / - | 10.45/11.75 | 0.01 | -0.01 | 3.27 | 0.01 | 36.10 |
| DD_DNA | bsc1 / - | 10.46/11.85 | -0.23 | 0.01 | 3.25 | -0.02 | 35.88 |
| DD_DNA | Tumuc1 / - | 9.59/10.99 | 0.05 | 0.00 | 3.29 | -0.01 | 36.78 |
| DD_DNA | DESRES / - | 11.68/12.92 | -0.58 | 0.00 | 3.24 | 0.00 | 34.19 |
| DD_DNA | CHARMM36 / - | 13.28/13.78 | 0.20 | 0.04 | 3.26 | -0.01 | 35.86 |
| DD_RNA | - / OL3 | 17.29/17.19 | -1.56 | 0.00 | 2.63 | 0.01 | 32.71 |
| DD_RNA | - / ROC-RNA | 16.75/16.87 | -1.65 | 0.04 | 2.76 | -0.05 | 32.31 |
| DD_RNA | - / DESRES | 16.76/16.79 | -2.03 | -0.01 | 2.72 | 0.02 | 30.01 |
| DD_RNA | - / CHARMM36 | 17.37/17.31 | -1.60 | 0.08 | 2.52 | -0.09 | 32.24 |
| DD_hybrid | OL21 / OL3 | 14.91/15.19 | -0.96 | 2.25 | 2.93 | -3.77 | 33.13 |
| DD_hybrid[a] | OL21 / OL3 | 17.24/16.98 | -1.37 | -0.36 | 2.72 | 0.64 | 32.95 |
| DD_hybrid | OL21 / OL3 (OPC water) | 15.00/15.12 | -1.05 | 2.16 | 2.94 | -3.66 | 32.97 |
| DD_hybrid | OL15 / OL3 | 14.94/15.19 | -0.96 | 2.18 | 2.95 | -3.69 | 33.34 |
| DD_hybrid[b] | OL15 / OL3 | 14.97/14.99 | -0.92 | 2.06 | 2.97 | -3.51 | 33.53 |
| DD_hybrid[c] | OL15 / OL3 | 15.29/15.36 | -1.18 | 2.01 | 2.91 | -3.60 | 32.72 |
| DD_hybrid | bsc1 / OL3 | 15.04/15.18 | -1.08 | 2.36 | 2.94 | -4.07 | 33.43 |
| DD_hybrid | bsc1 / OL3 (OPC water) | 14.97/15.13 | -1.16 | 2.32 | 2.94 | -4.03 | 33.01 |

| | | | | | | | |
|---|---|---|---|---|---|---|---|
| DD_hybrid | OL21 / ROC-RNA | 14.35/14.93 | -0.91 | 2.19 | 3.02 | -3.91 | 33.51 |
| DD_hybrid | Tumuc1 / OL3 | 14.84/15.17 | -1.18 | 2.63 | 2.96 | -4.75 | 33.00 |
| DD_hybrid | Tumuc1 / ROC-RNA | 14.26/14.74 | -1.01 | 2.49 | 3.05 | -4.55 | 33.42 |
| DD_hybrid | DESRES / DESRES | 15.46/15.46 | -1.51 | 2.15 | 2.91 | -4.14 | 31.22 |
| DD_hybrid | CHARMM36 / CHARMM36 | 16.76/16.81 | -1.17 | 2.14 | 2.82 | -3.56 | 33.87 |
| DD_hybrid[d] | CHARMM36 / CHARMM36 | 16.83/16.52 | -1.21 | 2.25 | 2.81 | -3.78 | 33.73 |
| PPT | *X-ray structure*[e] | 17.73 | -1.45 | -1.87 | 2.92 | 2.46 | 33.83 |
| PPT | OL21 / OL3 | 14.29 | -1.06 | 0.81 | 3.12 | -1.48 | 32.78 |
| PPT | OL15 / OL3 | 14,50 | -1.12 | 0.94 | 3.09 | -1.66 | 32.49 |
| PPT | bsc1 / OL3 | 14.41 | -1.22 | 1.01 | 3.10 | -1.79 | 32.40 |
| PPT | OL21 / ROC-RNA | 13.05 | -0.77 | 0.48 | 3.22 | -0.89 | 33.85 |
| PPT | Tumuc1 / OL3 | 14.12 | -1.30 | 1.32 | 3.26 | -2.39 | 32.63 |
| PPT | DESRES / DESRES | 15.03 | -1.36 | 1.17 | 3.02 | -2.20 | 31.38 |
| PPT | CHARMM36 / CHARMM36 | 16.86 | -1.53 | -0.35 | 2.82 | 0.72 | 31.81 |
| **Polarizable force fields** | | | | | | | |
| DD_DNA | DRUDE / - | 12.48/13.18 | -0.32 | -0.01 | 3.28 | 0.02 | 37.98 |
| DD_DNA | AMOEBA / - | 9.88/10.52 | -0.20 | -0.05 | 3.21 | 0.07 | 35.91 |
| DD_RNA | - / DRUDE | 17.27/17.32 | -1.26 | 0.00 | 2.91 | 0.00 | 34.59 |
| DD_RNA | - / AMOEBA | 16.88/16.95 | -2.26 | 0.01 | 2.80 | -0.03 | 30.83 |
| DD_hybrid | DRUDE / DRUDE | 15.49/15.96 | -0.98 | 2.94 | 3.13 | -4.69 | 35.38 |
| DD_hybrid | AMOEBA / AMOEBA | 16.77/16.14 | -2.08 | 0.69 | 2.83 | -1.35 | 30.44 |
| PPT | DRUDE / DRUDE | 14.67 | -1.12 | 1.50 | 3.43 | -2.63 | 34.27 |
| PPT | AMOEBA / AMOEBA | 15.11 | -1.54 | 0.81 | 3.11 | -1.62 | 31.85 |

[a]The DNA nucleotide puckers were restrained to C3′-endo region (pseudorotation values of -10° to 40°) by five flat-well dihedral restraint potential functions applied on each 2-deoxyribose (see Supporting Information).

[b]The initial structure of the hybrid corresponded to the B-form helix (see Methods).

[c]The $\chi$ dihedral potentials of the OL15 DNA *ff* were replaced with the RNA $\chi$ dihedral potentials from OL3 *ff*. See Ref. (1) for more details.

[d]2 kcal/mol stabilizing sHBfix potential was applied on every base pairing H-bond.

[e]Average values observed in the PPT experimental structure (2). See Table S1 for values observed for other experimental structures of the hybrids.

### Supporting Information Figures

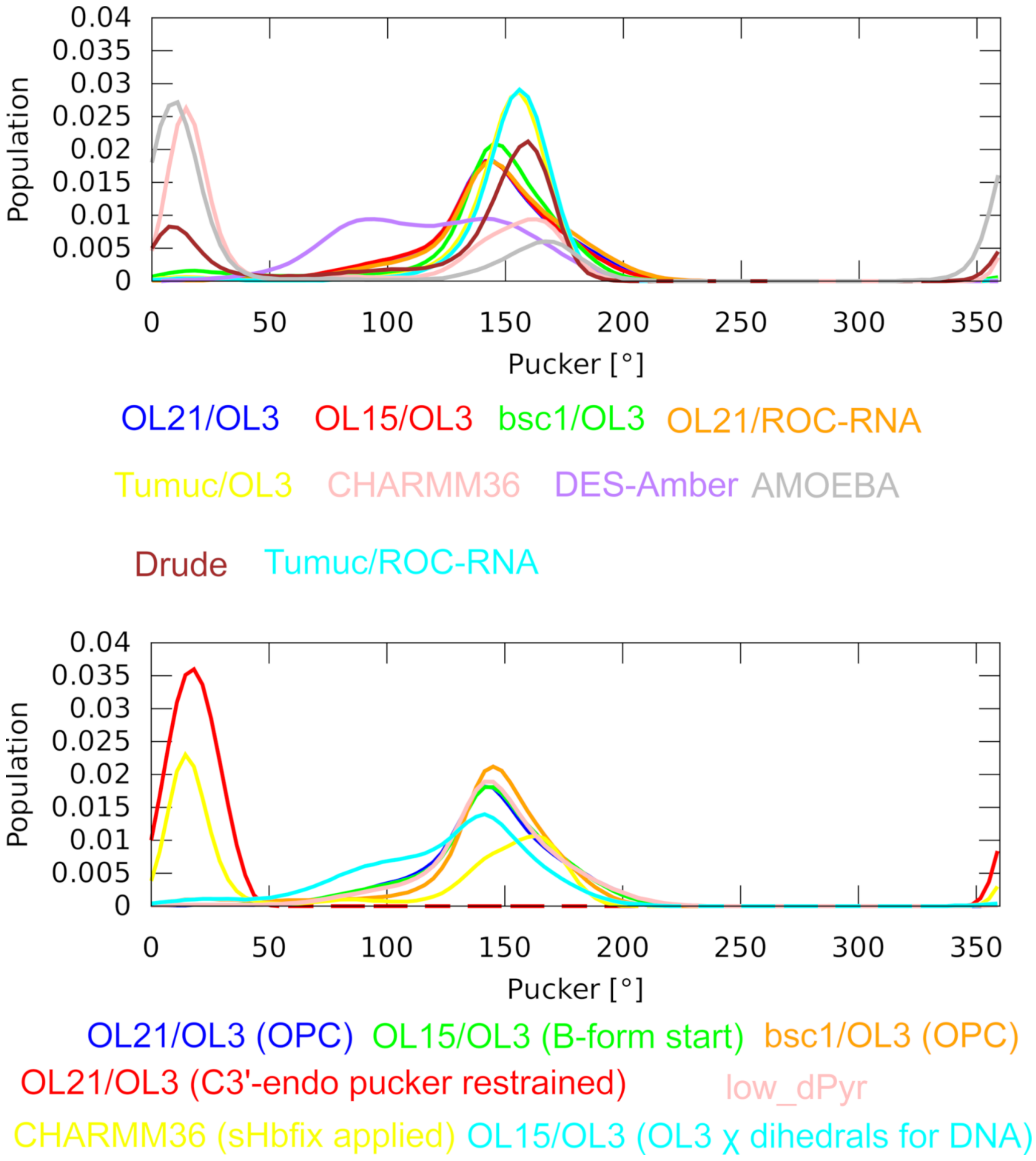


Figure S1. **Histograms of the DNA sugar puckers in MD simulations of the DD_hybrid structure and the low_dPyr structure.** Due to the sheer number of simulations, the datasets were separated into two graphs for the sake of clarity. The individual force-field combinations are color-coded for each graph according to the legend below the graph. For additional explanation of the data sets in the bottom graph, see the main text Methods and Table 1.

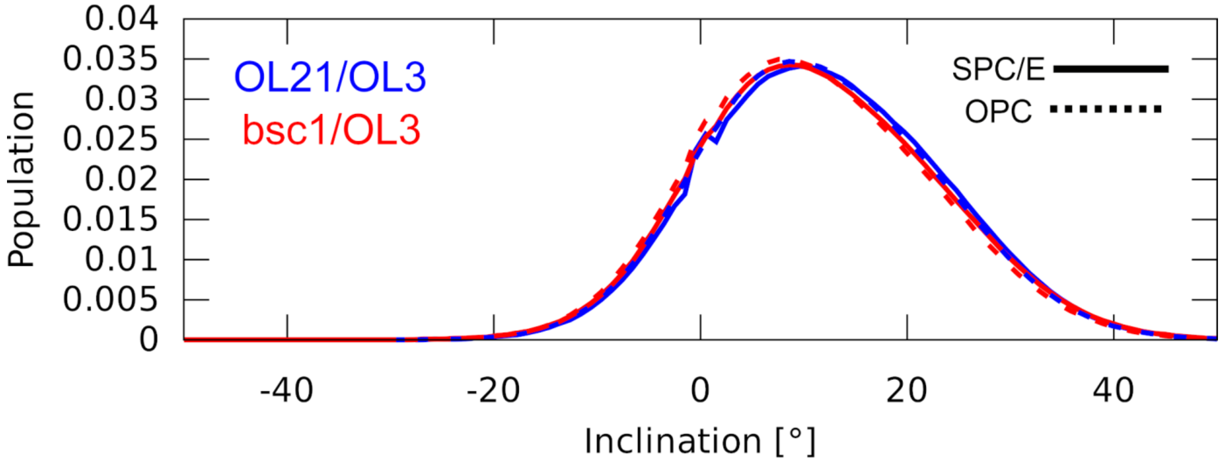

Figure S2. **Inclination of the DD_hybrid structure simulated using the SPC/E and OPC water models.** The individual force-field combinations are color-coded according to the legend. Full and dashed lines indicate SPC/E and OPC simulations, respectively.

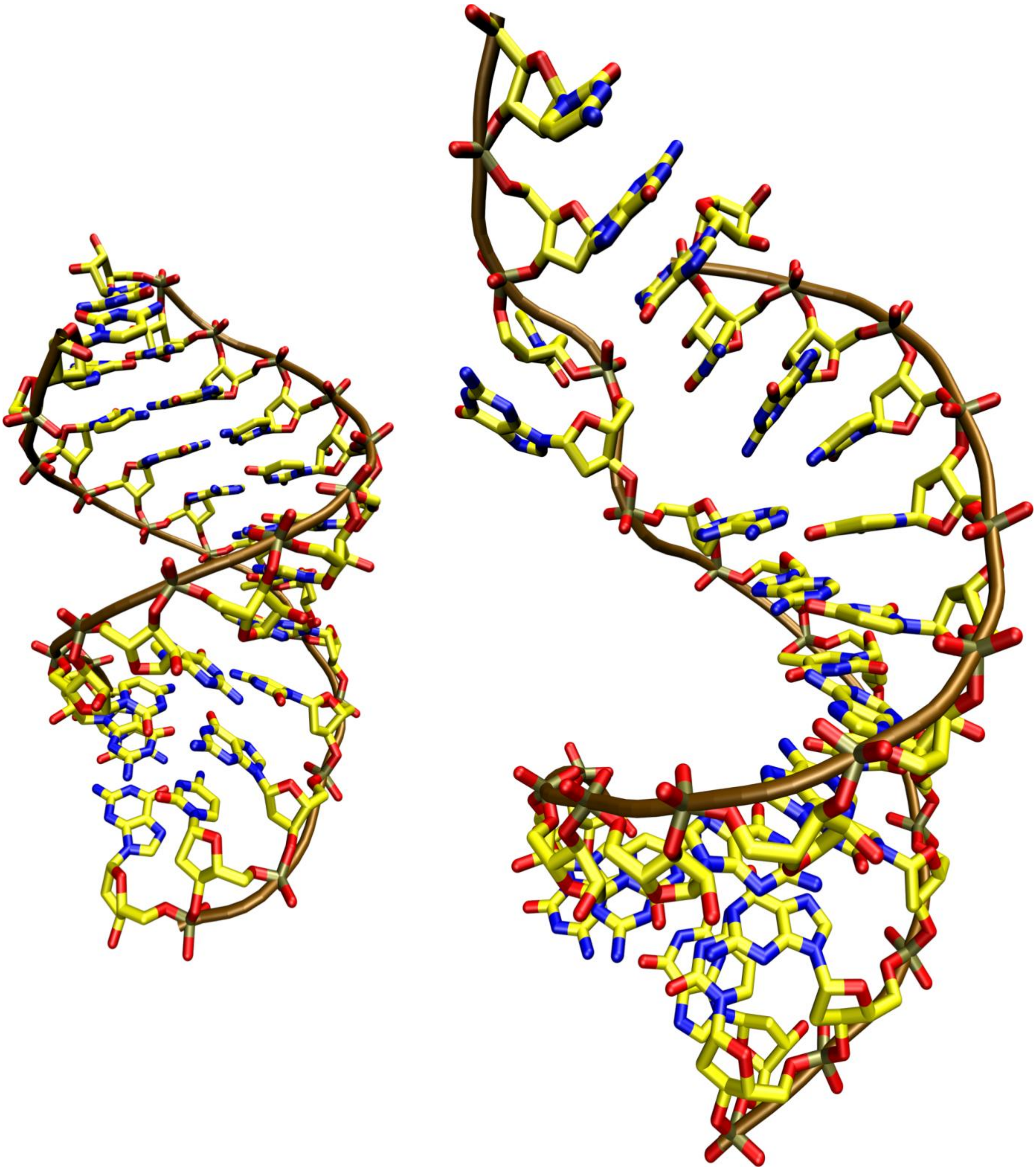

Figure S3. **Example of the extensive loss of base pairing sometimes observed in MD simulations using the CHARMM36 *ff*.** Two snapshots of the DD_hybrid structure are shown, a stable helix (left) and one with four consecutive GC base pairs disrupted in the upper part (right). The disruptions can be prevented by applying sHBfix on every base pair (see the main text).

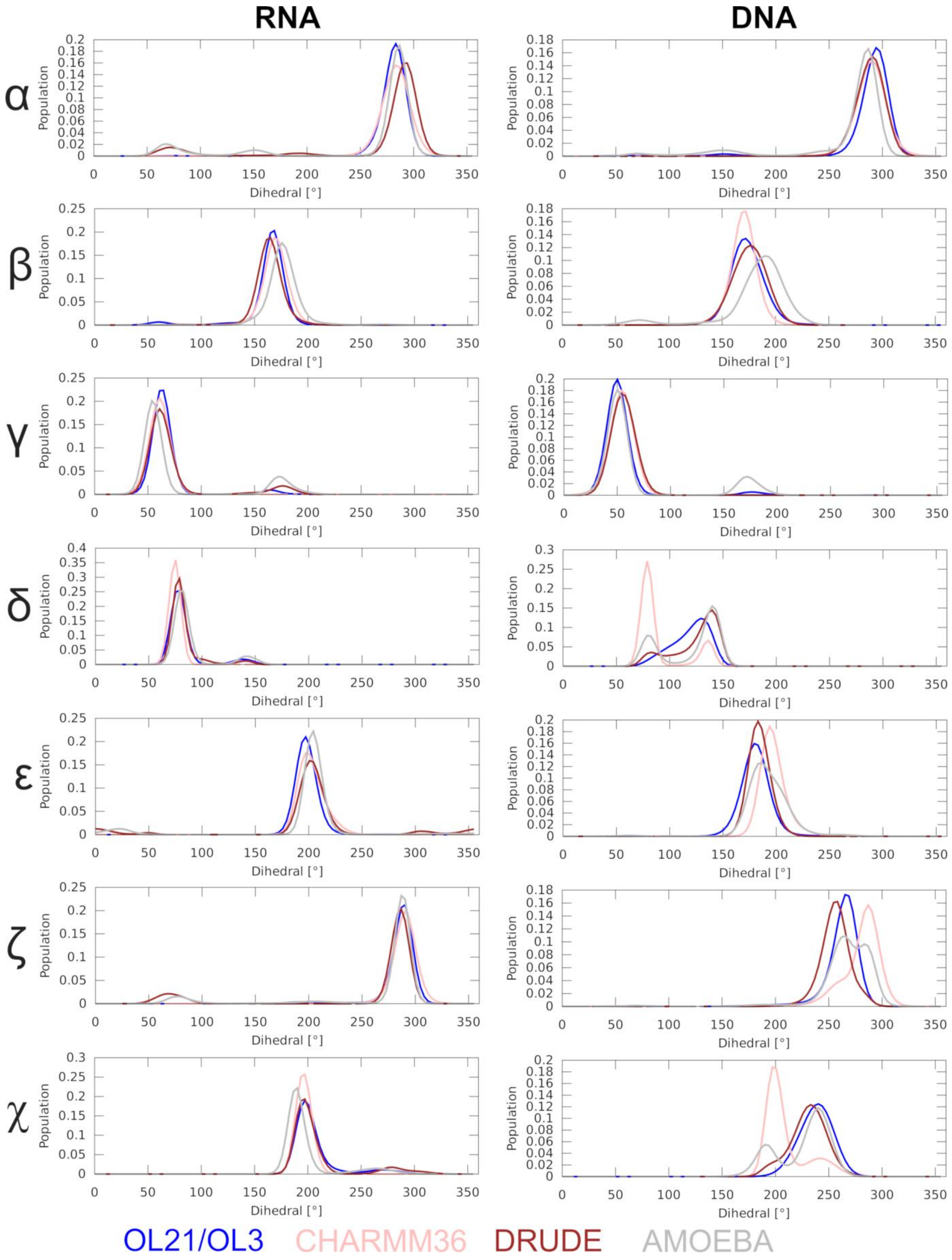


Figure S4. **Comparison of the backbone dihedral angles observed in MD simulations of the PPT hybrid structure using the polarizable and selected non-polarizable force fields.**

### Supporting Information References


1. Jurečka, P., Zgarbová, M., Černý, F. and Salomon, J. (2024) Multistate B- to A-Transition in Protein-DNA Binding – How Well is it Described by Current AMBER Force Fields? *J. Biomol. Struct. Dyn.*, 1-11.
2. Kopka, M.L., Lavelle, L., Han, G.W., Ng, H.-L. and Dickerson, R.E. (2003) An Unusual Sugar Conformation in the Structure of an RNA/DNA Decamer of the Polypurine Tract May Affect Recognition by RNase H. *J. Mol. Biol.*, **334**, 653-665.
